# Gesture profiles distinguish primary progressive aphasia variants

**DOI:** 10.1101/2023.01.19.524719

**Authors:** Haley C. Dresang, Rand Williamson, Hana Kim, Argye E. Hillis, Laurel J. Buxbaum

## Abstract

Primary progressive aphasia (PPA) is a neurodegenerative syndrome characterized by progressive language deficits. There are three main variants of PPA – semantic (svPPA), logopenic (lvPPA), and nonfluent (nfvPPA) – that can be challenging to distinguish. Limb praxis may also be affected in PPA, but it is unclear whether different variants of PPA are associated with differences in gesture production. Prior research with neurotypical individuals indicates that the left temporal lobe is a critical locus of manipulable object and hand posture representations. Moreover, when imitating gestures, individuals whose strokes include the left temporal lobe show reduced benefit of gesture meaning and disproportionate impairment in hand posture as compared to arm kinematics. We tested the hypothesis that svPPA – who typically exhibit primarily temporal lobe atrophy – would differentially show these expected patterns of gesture imitation performance. Nineteen participants with PPA completed meaningful and meaningless gesture imitation tasks, and performance was scored for hand posture and arm kinematics accuracy. Generalized logistic mixed-effect regression models controlling for dementia severity showed overall benefits from gesture meaning, and greater impairments in hand posture than arm kinematics. We also found that svPPA participants were the most impaired in gesture imitation overall. Critically, there was also a significant three-way interaction of group, meaning, and gesture component: only svPPA participants showed relative impairments of hand posture for meaningful gestures as well as meaningless gestures. Thus, unlike lvPPA and nfvPPA, the hand postures of svPPA failed to benefit from gesture meaning. This research extends prior findings on the role of the temporal lobe in hand posture representations associated with manipulable objects, and is the first to indicate that there may be distinct gesture imitation patterns as a function of PPA variant. Characterizing componential gesture deficits in PPA may help to inform differential diagnosis, compensatory communication strategies, and cognitive praxis models of PPA.

## 1. Introduction

Primary progressive aphasia (PPA) is a clinical syndrome caused by neurodegenerative disease such as frontotemporal dementia or Alzheimer’s disease that is characterized by progressive language deficits with relative sparing of other cognitive domains (Gorno-Tempini et al., 2011; Mesulam, 2003). However, it is also recognized that aspects of cognition and praxis may be deficient in PPA along with language (de la Sablonnière et al., 2021; Joshi et al., 2003; Yliranta & Jehkonen, 2020). There are three widely recognized clinical variants of PPA that are distinguished by their functional deficits and sites of neural atrophy. The semantic variant of PPA (svPPA) begins with bilateral anterior and inferior temporal lobe pathology that progresses to the posterior temporal lobes (Gorno-Tempini et al., 2011; Meeter et al., 2017). Patients with svPPA exhibit conceptual processing impairments and loss of semantic knowledge that is evident regardless of modality of input (e.g., visual versus verbal; Gorno-Tempini et al., 2011; Suárez-González et al., 2021). The nonfluent or agrammatic variant of PPA (nfvPPA) presents with grammatical deficits and/or apraxia of speech, along with progressive atrophy of left posterior fronto-insular regions (Gorno-Tempini et al., 2011). The logopenic variant of PPA (lvPPA) is associated with phonological speech errors, diminished short-term memory, impaired verbal repetition, and predominant left posterior perisylvian or parietal atrophy (Gorno-Tempini et al., 2011).

The differential diagnosis of PPA variants is challenging. Over 90 percent of individuals with frontotemporal dementia present with behavioral and linguistic symptoms (Harris et al., 2016), and most patients with Alzheimer’s disease pathology also have both linguistic and nonlinguistic cognitive deficits. Despite international consensus criteria to identify PPA subtypes, there is substantial overlap in the language characteristics of the 3 variants, vast variability in clinical presentation, and challenging speech and language features that can obscure differential diagnosis (Leyton & Hodges, 2014; Tippett, 2020). These challenges suggest that it would be useful to identify non-linguistic domains in which PPA variants may have different profiles. One possible domain is limb apraxia.

Limb apraxia – a disorder of skilled movements not attributable to sensory, motor, or language deficits – frequently co-occurs with PPA (Yliranta & Jehkonen, 2020; Buxbaum & Kalénine, 2021). Limb apraxia can affect skilled action production, imitation, recognition, and/ or tool use. The classical literature distinguishes 2 subtypes of limb apraxia: (1) ideational apraxia, characterized by deficit or loss of action concepts including sequential and spatial procedures for complex, multi-step actions; (2) ideomotor apraxia, characterized by difficulty retrieving kinematic patterns and integrating these patterns into a movement plan (DeRenzi & Lucchelli, 1988; Liepmann, 1988). However, more recent investigations from our group and others indicate that the classical dichotomy has been difficult to validate. Hereafter, we use the term “apraxia” broadly to encompass both subtypes (see Buxbaum & Kalénine, 2021 for discussion).

Gesture pantomime imitation is a common assessment used to identify and characterize limb apraxia. Cognitive praxis models posit two neurocognitive routes enabling gesture imitation: a direct route, which translates the visual input—a gesture performed by another individual – into a motor plan via intermediate body representations; and an indirect route, which additionally incorporates access to conceptual and spatiotemporal representations of familiar actions (Achilles et al., 2019). The direct route is most clearly assessed via imitation of meaningless or novel gestures, whereas the indirect route can be assessed with measures of action knowledge, recognition, production of gestures to the sight of tools, and imitation of meaningful, familiar actions (Buxbaum & Kalénine, 2021; Tessari et al., 2007, 2021). Just as is the case with word versus non-word repetition in the language domain, gesture imitation in neurotypical individuals is typically more accurate when gestures are meaningful; thus, meaningful content confers a processing advantage.

There is also evidence that semantic meaning provides a benefit for gesture production in individuals with stroke. In particular, individuals with chronic left-hemisphere stroke more accurately imitate meaningful than meaningless gestures (Bartolo et al., 2001; Goldenberg & Hagmann, 1997; Tessari et al., 2007). We have demonstrated that adults with left-hemisphere stroke perform better on both labeled and unlabeled meaningful compared to meaningless gestures, consistent with the involvement of the indirect semantic route in meaningful gesture imitation (Dresang et al., in press). Conversely, reduced benefit from meaning in imitation points to compromised integrity of this semantic processing route and is associated with temporal lobe damage. Moreover, deficits in meaningful but not meaningless gesture imitation have been associated with naming and word repetition impairments following stroke (Mengotti et al., 2013).Together, these data lead to the prediction that compared to individuals with relatively intact semantics (lvPPA, nfvPPA), individuals with semantic impairments – such as svPPA – may show reduced semantic benefits on gesture production ability.

Prior research indicates that limb apraxia may be characterized by deficits in two major components of gesture: (1) hand posture, which includes the shape and movements of the fingers, hand, and wrist (e.g., the movements of the fingers/hand/ wrist needed to turn a screwdriver); and (2) arm kinematics, or the amplitude and timing of the arm movements (e.g., the characteristic “swinging” movement of a hammer; Jax & Buxbaum, 2013).^1^ Although few studies have assessed these components, there is evidence that they may dissociate following left hemisphere stroke.

Our prior research indicates that hand posture errors may be observed in imitation of both meaningless gestures and meaningful tool-specific pantomime gestures, as well as in gestures produced in response to viewed tools (Buxbaum et al., 2014). Hand posture errors are also associated with deficits in knowledge of actions and manipulable objects (i.e., action semantics), as well as with left posterior temporal lesions (Dresang et al., in press; Tarhan et al., 2015). These data are consistent with numerous functional neuroimaging studies in neurotypical adults that identify posterior temporal regions associated with both hand and manipulable object representations (Bergström et al., 2021; Bracci et al., 2016, 2018), and with studies that characterize posterior temporal cortex as the brain’s major hub for tool semantics (van Elk et al., 2014). More broadly, both the middle temporal gyrus and anterior temporal lobe have been suggested to mediate tool and hand representations for tool-use actions (Knights et al., 2022; Lesourd et al., 2021). In contrast, kinematics errors are associated with meaningless gesture imitation and with damage to pre/postcentral gyri and inferior parietal lobe regions, including supramarginal gyrus (Buxbaum et al., 2014; Watson & Buxbaum, 2015). The inferior parietal lobe has also been associated with spatiotemporal aspects of action prediction and planning (de Wit & Buxbaum, 2017; Urgen & Saygin, 2020). Within the frontal lobe, prefrontal and premotor cortices are essential for context-, goal-, and task-dependent action selection (e.g., Badre, 2008).

To our knowledge, hand posture and arm kinematic gesture components have never been assessed in PPA. We reasoned that characterizing componential gesture deficits in PPA might help to inform differential diagnosis of PPA variants and other forms of dementia, including Alzheimer’s dementia. Furthermore, praxis and gesture are vital for communication, and characterizing the components of limb apraxia affected in PPA could inform speech-language therapy and compensatory communication strategies. Finally, these data would be useful in assessing the universality of cognitive praxis models, which have predominantly been informed by evidence from neurotypical adults and adults with left-hemisphere stroke.

A recent systematic scoping review compiled the evidence regarding limb and face apraxia in PPA (Yliranta & Jehkonen, 2020). The authors reported that limb apraxia has been identified in all 3 variants of PPA, with an overlapping range of apraxia frequency reported in each variant: 40%-58% in nfvPPA, 36%-67% in lvPPA, and 69%-90% in svPPA. Therefore, the presence or frequency of apraxia is not sufficient to differentiate between variants of PPA. However, svPPA was uniquely characterized by deficits on tasks that required action semantic knowledge (e.g., action matching tasks). Individuals with svPPA also uniquely showed increased production of content errors (i.e., perplexity or unrecognizable gestures) with less substantial deficits in the spatiotemporal aspects of gesture (Yliranta & Jehkonen, 2020).

Based on the available evidence from individuals with stroke and neurotypical individuals, we tested 3 sets of predictions. First, we tested the prediction that as a group, individuals with PPA would more accurately imitate meaningful than meaningless gestures. Second, we predicted that kinematic movements would be imitated more accurately than hand postures, as has been shown in neurotypical controls and individuals with stroke. Finally, we predicted that individuals with svPPA would exhibit a different pattern of gesture deficits than lvPPA or nfvPPA, motivated by previously demonstrated close associations between gesture hand posture and semantic deficits, as well as by known differences in the neuroanatomic loci of atrophy in each variant of PPA. Specifically, we predicted that: (1) Relative to kinematics, hand posture deficits would be more pronounced in svPPA than in lvPPA or nfvPPA; and (2) Hand posture during gesture imitation in participants with svPPA would not benefit from gesture meaning, as it would in participants with lvPPA and nfvPPA.

## 2. Methods

### 2.1. Participants

Participants were 19 adults with diagnosed with PPA by meeting consensus criteria for one of the variants of PPA (Gorno-Tempini et al., 2011), based on performance on a battery of language and cognitive tests and imaging. The battery covered both nonlinguistic domains of cognition and the main language domains (naming, word and sentence comprehension, repetition, syntactic processing, reading, spelling, and non-verbal semantics). The following tests were administered: Boston Naming Test (Kaplan et al., 1983) or 30 item short form of the Boston Naming Test (Mack et al., 1992); Hopkins Assessment of Naming Actions (Breining et al., 2022); National Alzheimer’s Coordinating Center (NACC) Frontotemporal Lobar Degeneration Module (Gefen et al., 2020) subtests of Figure Copy, Recall, and Recognition, Word Fluency, Sentence Repetition, Sentence Reading subtest, and Word-picture matching; Pyramids and Palm Trees test short version (Breinig et al., 2015); Kissing and Dancing Test short version (unpublished version of Mansur et al., 1980); and Word-Picture Verification with 30 objects and 30 actions (with semantic foils, phonological foils, and targets; Kay et al., 2009). Using neurological exam, history, the language and cognitive battery, and imaging, four participants had semantic variant PPA (svPPA), five had nonfluent variant PPA (nfvPPA), and ten had logopenic variant PPA (lvPPA). All participants had normal or corrected-to-normal vision, no hearing loss, and no history of other neurologic disease. To further characterize the degree of severity, dementia severity (Dementia Rating Scale: Porto et al., 2007) was evaluated in all participants.^2^ In compliance with the guidelines of the Institutional Review Board of Johns Hopkins University, all participants provided informed consent. Demographic information is reported in Table 1.

**Table 1.** Participant demographic characteristics.

| Participant ID | Age | Months Post-Diagnosis | Dementia Severity | Education | Sex | Handedness |
| --- | --- | --- | --- | --- | --- | --- |
| svPPA01 | 71 | 37 | 5.5 | 16 | F | R |
| svPPA02 | 69 | 7 | 7.5 | 12 | F | R |
| svPPA03 | 67 | 50 | 6 | 16 | F | R |
| svPPA04 | 60 | 39 | 6 | 16 | F | R |
| nfvPPA01 | 85 | 6 | 1.5 | n/a | F | n/a |
| nfvPPA02 | 63 | 34 | 1.5 | 18 | F | R |
| nfvPPA03 | 73 | n/a | n/a | n/a | M | R |
| nfvPPA04 | 62 | n/a | 0.5 | n/a | M | R |
| nfvPPA05 | 74 | 9 | 8 | 18 | M | R |
| lvPPA01 | 54 | 25 | 11 | 16 | F | n/a |
| lvPPA02 | 73 | 12 | 1.5 | 18+ | M | L |
| lvPPA03 | 75 | 1 | 1.5 | n/a | F | n/a |
| lvPPA04 | 72 | 3 | 5.5 | n/a | M | n/a |
| lvPPA05 | 82 | 2 | 4.5 | 14 | F | R |
| lvPPA06 | 62 | 9 | 0.5 | 18 | M | R |
| lvPPA07 | 65 | n/a | 0 | 16 | F | R |
| lvPPA08 | 63 | n/a | 0 | 16 | M | L |
| lvPPA09 | 69 | n/a | 0 | 16 | M | R |
| lvPPA10 | 73 | n/a | 1 | 16 | M | R |

### 2.2. Materials and procedures

#### Gesture imitation experimental task

Participants completed two video-based gesture imitation tasks: (1) meaningless gestures and (2) meaningful transitive gestures (e.g., pantomimed hammering). Parallel to previous protocols, participants were instructed to use their left hand. For each task, videos showed an experimenter performing 16 gestures with the right hand. Participants were instructed to imitate each gesture as if looking in a mirror. Each gesture was presented two times; participants were instructed to watch the first presentation of the gesture and to begin imitating along with the model when they heard a tone that played at the start of the second presentation. The duration of each gesture video was 4-5 seconds. The meaningless and meaningful gestures were matched for similar spatiomotor characteristics (e.g., plane of movement, joints moved, hand posture; as described in Buxbaum et al., 2014). Meaningful gestures were recognized reliably by ten right-handed control participants (M = 97.5% accuracy, SD = 3.8%, range = 91.7 – 100%; as described in Tarhan et al., 2015).

#### Gesture coding

Each participant’s gesture production was video recorded and scored offline by a trained coder who had achieved inter-rater reliability (Cohen’s kappa) of >85% with other raters in the lab. Each trial was assigned a score of 0 (incorrect) or 1 (correct) for each gesture component – hand posture and kinematics – following established procedures implemented in numerous studies (see Buxbaum et al., 2000 for details). See Table 2 for a summary of participant performance on each component per gesture task.

**Table 2.**
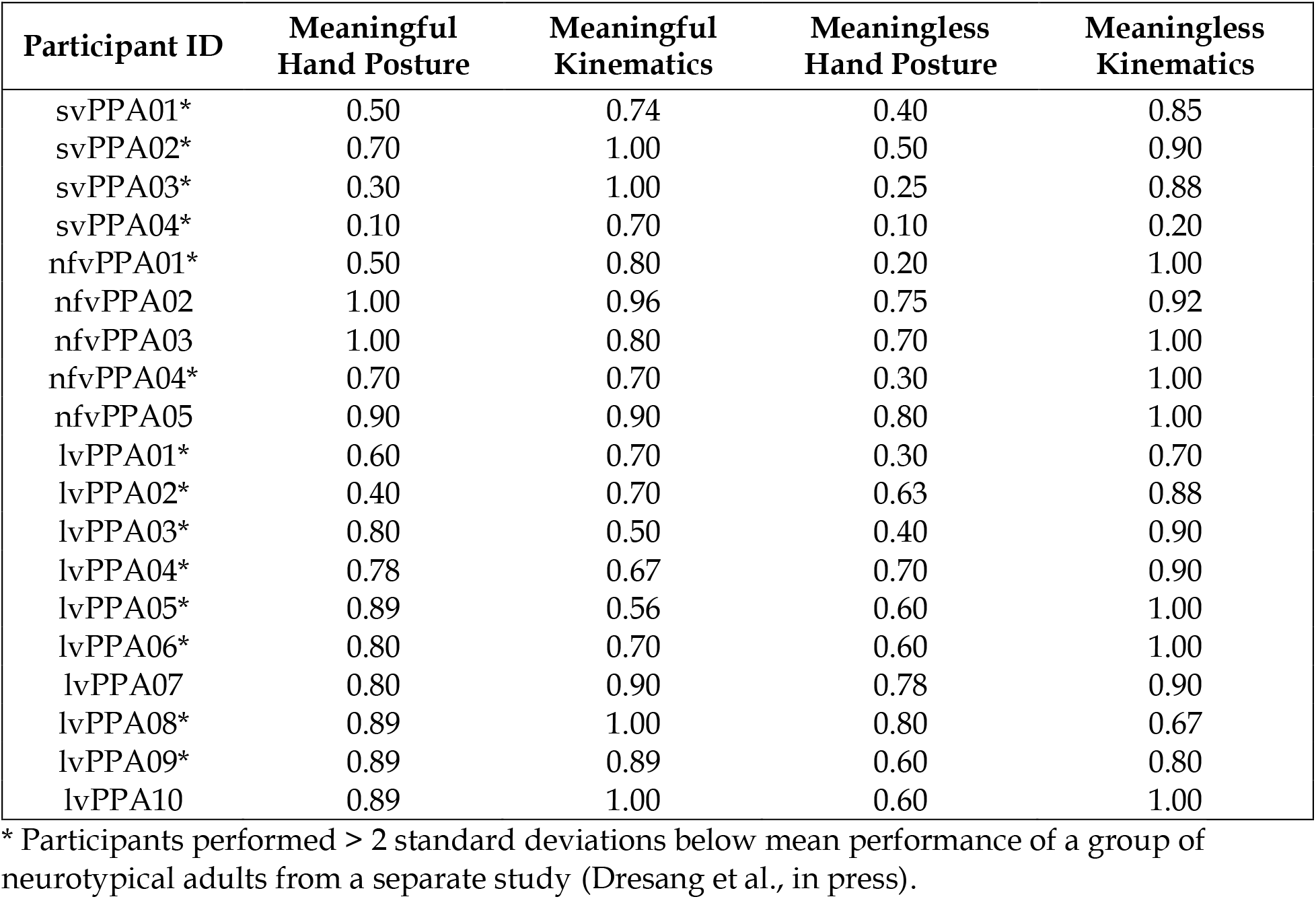
Participant mean performance across tasks.

| Participant ID | Meaningful Hand Posture | Meaningful Kinematics | Meaningless Hand Posture | Meaningless Kinematics |
| --- | --- | --- | --- | --- |
| svPPA01* | 0.50 | 0.74 | 0.40 | 0.85 |
| svPPA02* | 0.70 | 1.00 | 0.50 | 0.90 |
| svPPA03* | 0.30 | 1.00 | 0.25 | 0.88 |
| svPPA04* | 0.10 | 0.70 | 0.10 | 0.20 |
| nfvPPA01* | 0.50 | 0.80 | 0.20 | 1.00 |
| nfvPPA02 | 1.00 | 0.96 | 0.75 | 0.92 |
| nfvPPA03 | 1.00 | 0.80 | 0.70 | 1.00 |
| nfvPPA04* | 0.70 | 0.70 | 0.30 | 1.00 |
| nfvPPA05 | 0.90 | 0.90 | 0.80 | 1.00 |
| lvPPA01* | 0.60 | 0.70 | 0.30 | 0.70 |
| lvPPA02* | 0.40 | 0.70 | 0.63 | 0.88 |
| lvPPA03* | 0.80 | 0.50 | 0.40 | 0.90 |
| lvPPA04* | 0.78 | 0.67 | 0.70 | 0.90 |
| lvPPA05* | 0.89 | 0.56 | 0.60 | 1.00 |
| lvPPA06* | 0.80 | 0.70 | 0.60 | 1.00 |
| lvPPA07 | 0.80 | 0.90 | 0.78 | 0.90 |
| lvPPA08* | 0.89 | 1.00 | 0.80 | 0.67 |
| lvPPA09* | 0.89 | 0.89 | 0.60 | 0.80 |
| lvPPA10 | 0.89 | 1.00 | 0.60 | 1.00 |

\* Participants performed > 2 standard deviations below mean performance of a group of neurotypical adults from a separate study (Dresang et al., in press).

### 2.3. Data analysis

Data were analyzed using generalized logistic mixed-effect regression models in the R statistical programming environment with the lme4 package (Bates et al., 2015). Trial-level gesture imitation accuracy was the dependent variable. Fixed effects included a three-way interaction between PPA variant (svPPA, nfvPPA, lvPPA), task condition (meaningful, meaningless gestures), and gesture component (hand posture, kinematics), controlling for dementia severity (Dementia Rating Scale: Porto et al., 2007). Random intercepts were included for subject and item. We assessed all mixed-effect models by mapping the log likelihood ratio of full and reduced models using a chi square distribution. In addition, we removed interaction terms and assessed the full model when testing for significant main effects. We used an alpha threshold of 0.05 to determine statistical significance. For planned post hoc tests, we used the emmeans package and the Tukey adjustment to correct for multiple comparisons (Lenth et al., 2022).

## 3. Results

See Table 2 for a summary of participant performance on each component per gesture task. Fourteen of the 19 participants performed more than two standard deviations below the mean of a group of neurotypical subjects who were reported in a previous paper (Dresang et al., in press). By this measure, 100% of svPPA, 80% of lvPPA, and 40% of nfvPPA may broadly be classified as apraxic. Dementia severity was not correlated with performance on any gesture task (all p-values > 0.0125, Bonferroni corrected significance value).

There were significant main effects of each predictor after controlling for dementia severity. In particular, a main effect of PPA variant suggested that svPPA were more impaired in gesture imitation than the other PPA variants (χ^2^ = 15.56, p = 0.04). A main effect of gesture component indicated that hand posture was, on average, more impaired than kinematics (χ2 = 73.97, p < 0.001). A main effect of task condition showed that meaningless gestures were, on average, more impaired than meaningful gesture imitation (χ2 = 21.00, p = 0.002). Moreover, there was a significant three-way interaction between PPA variant, gesture component, and task condition (χ2 = 24.95, p < 0.001; Figure 1). Pairwise comparisons with Tukey corrections revealed that at the group level, hand posture was more impaired than kinematics for all PPA variants in the meaningless condition (Table 3). In contrast, only svPPA participants showed relative impairment of hand posture compared to kinematics for meaningful gestures (β = 1.98, p = 0.007). At the individual level, for 7 participants (4 lvPPA, 3 nfvPPA), kinematics was more impaired than hand posture for meaningful gestures. Importantly, none of the participants showing this superiority of hand posture over kinematics components were svPPA.

**Figure 1.**
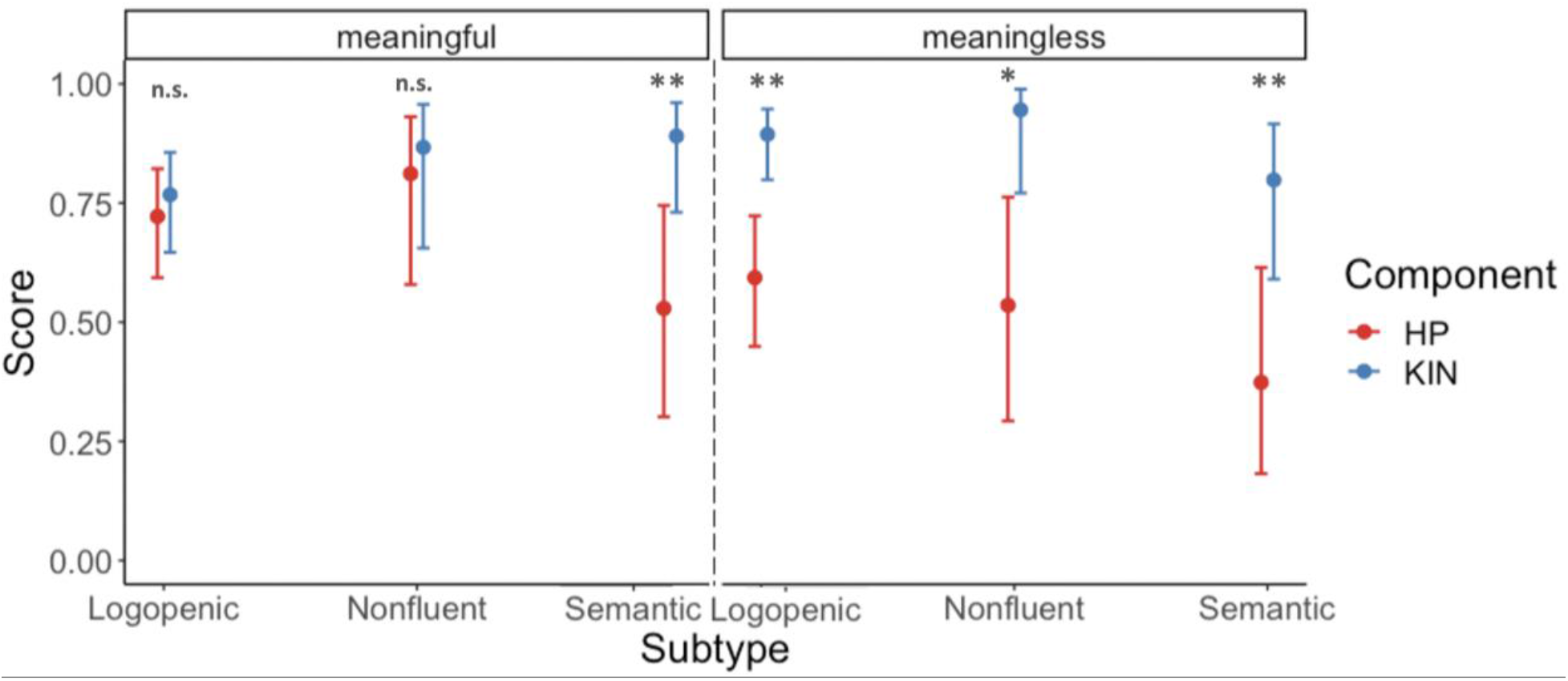
Three-way interaction between PPA variant x gesture component x gesture meaning interaction. *Note:* HP: Hand posture component score, KIN: Kinematic component score.

**Table 3.** Pairwise comparisons (with Tukey corrections) of the PPA variant x gesture component x gesture meaning interaction.

| PPA Variant | Meaningful Kinematics-Hand Posture Contrast |  |  |  | Meaningless Kinematics-Hand Posture Contrast |  |  |  |
| --- | --- | --- | --- | --- | --- | --- | --- | --- |
|  | estimate | SE | z ratio | p value | estimate | SE | z ratio | p value |
| svPPA | 1.98 | 0.52 | 3.83 | 0.007 | 1.89 | 0.48 | 3.95 | 0.005 |
| lvPPA | 0.24 | 0.31 | 0.78 | 0.99 | 1.76 | 0.40 | 4.44 | 0.001 |
| nvPPA | 0.41 | 0.64 | 0.64 | 1.00 | 2.71 | 0.81 | 3.34 | 0.040 |

## 4. Discussion

Hand posture and arm kinematics are two major components of gesture that can be impaired in limb apraxia and are associated with different behavioral profiles and lesion loci in individuals who have experienced stroke. These gesture components have never been assessed in PPA, and cognitive praxis models have predominantly been informed by evidence from neurotypical adults and adults with left-hemisphere stroke. Prior work in neurotypicals and stroke survivors demonstrates that semantic meaning confers benefits for gesture production (Dresang et al., in press; Goldenberg & Hagmann, 1997; Tessari et al., 2007). However, the facilitative effect of meaning has not been previously examined in PPA praxis and is likely to differ based on PPA variant. Characterizing gesture component deficits and differential benefits of semantic meaning in PPA may be instrumental in informing differential diagnosis of PPA variants and the utility of gesture-based compensatory communication strategies for PPA. Therefore, the purpose of this study was to evaluate hand posture and kinematic components of meaningful and meaningless gestures across all three variants of PPA.

First, we tested the prediction that as a group, individuals with PPA would show effects of gesture component and gesture meaning, parallel to what has been reported in neurotypical adults and individuals living with stroke. In line with our predictions, we found effects of component and meaning on gesture imitation. In particular, after controlling for overall dementia severity, our sample of PPA showed an average benefit from semantic information, such that meaningful gestures were produced more accurately than meaningless gestures. Similarly, our sample of PPA showed significantly greater impairments in hand posture than kinematic gesture components. These findings are broadly consistent with evidence from stroke patients (see Buxbaum & Kalénine, 2021 for review) and critically extend scientific knowledge of the nature of limb apraxia deficits in PPA. Although the current study did not evaluate neurotypical adults, published neurotypical data from the same gesture assessments (Dresang et al., in press), showed that neurotypical adults show better performance on limb praxis than many PPA participants (and all svPPA). Our data provide preliminary evidence that individuals with PPA can demonstrate signs of limb apraxia during meaningful gesture imitation, meaningless gesture imitation, or both. Future work should consider the effects of gesture deficits in individuals with PPA (Joshi et al., 2003).

Second, we tested the prediction that svPPA would exhibit a different pattern of gesture deficits than lvPPA or nfvPPA. Specifically, we predicted that relative to arm kinematics, hand posture deficits would be more pronounced in svPPA than lvPPA or nfvPPA participants. We also predicted that participants with svPPA would show reduced benefits from gesture meaning on hand posture, as compared to lvPPA and nfvPPA participants. Overall, svPPA were the most impaired in gesture imitation when we controlled for dementia severity. Moreover, there was a significant three-way interaction between gesture components, meaning, and PPA variant. For meaningful gestures, svPPA were the only group who were more impaired in imitating hand postures relative to kinematic gesture components; that is, unlike the other PPA variants, their hand postures failed to benefit from gesture meaning. These findings suggest that compared to other PPA variants, individuals with svPPA have a marked impairment in imitating the hand posture component of meaningful gestures, consistent with prior findings that action (e.g., object manipulation) knowledge is impaired in patients with semantic dementia (Meligne et al., 2011). Thus, hand posture deficits in meaningful gestures may be a useful behavioral marker to help distinguish svPPA from other PPA variants. From a cognitive neuropsychological perspective, this pattern of results is consistent with prior evidence that hand posture gesture errors are associated with deficits in semantic knowledge of actions and manipulable objects, as well as with temporal lobe lesions (Buxbaum et al., 2014; Dresang et al., in press; Tarhan et al., 2015).

In contrast to the current findings, a recent study suggested that nonverbal action knowledge may be a relative strength – in comparison to nonverbal object knowledge – in individuals with svPPA (Auclair-Ouellet et al., 2020). However, these conclusions were based on semantic picture matching and naming assessments and not gesture production tasks. Hand posture during meaningful gesture imitation requires knowledge of both object and action semantics, including information such as the relevant affordances of the object that is associated with the viewed pantomime gesture, and the context of the object’s manipulation. During gesture imitation tasks, individuals must activate the indirect semantic route, entailing access to stored object and action knowledge, to enable meaningful gestures to be imitated more accurately than meaningless gestures (Buxbaum et al., 2005; Dresang et al., in press). Accurate meaningful hand posture imitation requires knowledge of the shape and size of affordances of the object associated with the viewed gesture, as well as knowledge of the hand and finger movements needed to perform a functional action with that object. For example, upon viewing a cutting gesture, participants must integrate knowledge of scissors and their affordances, as well as the precise hand posture required for a cutting motion. Picture matching and naming assessments may not be sensitive to this same depth of object and action knowledge.

Lesion studies from individuals with stroke have shown that the left posterior temporal lobe is critical for knowledge of manipulable objects and their associated hand postures (Bracci et al., 2018; Dresang et al., in press; Tarhan et al., 2015). This has also been supported by functional neuroimaging evidence that associates posterior temporal lobe activation with hand posture and manipulable object representations (Bergström et al., 2021; Bracci et al., 2016, 2018). The anterior temporal lobe, too, may be critical in combining semantic and sensorimotor information relevant to tool manipulation and gesture production (Knights et al., 2022; Lesourd et al., 2021; Peelen & Caramazza, 2012). Given that svPPA have pathology affecting the anterior and, as the disease progresses, more posterior temporal lobes (Gorno-Tempini et al., 2011; Meeter et al., 2017), both regions may contribute to the marked deficits in hand posture components of meaningful gestures observed in the svPPA group.

The present results suggest that the assessment of limb apraxia in relation to semantic knowledge in PPA should be further investigated. The findings represent a novel extension of prior research on praxis in PPA (Joshi et al., 2003), but there were several limitations. Our study involved a convenience data sample, and future work may benefit from a larger sample size with a robust distribution of PPA variants, additional measures of aphasia, praxis, and semantic processing, as well as concurrent neuroimaging to characterize location and degree of atrophy associated with praxis deficits in individual participants.

## 5. Conclusion

This study is the first, to our knowledge, to replicate prior findings from stroke and neurotypical individuals that meaning generally confers a benefit for gesture imitation in individuals with PPA. In addition, we demonstrated that hand posture was more impaired than arm kinematics components for meaningless gesture imitation across all PPA variants, consistent with evidence that hand posture is the most sensitive measure of limb apraxia in patients with left-hemisphere stroke (Jax & Buxbaum, 2013). However, controlling for dementia severity, individuals with svPPA were the most severely apraxic overall. Importantly, svPPA was the only variant that also showed relative deficits in hand posture for meaningful gestures, and no participants with svPPA performed better on hand posture than kinematic gesture components. These findings suggest that degraded semantic representations or retrieval processes in svPPA are tightly linked to deficient hand posture accuracy, possibly mediated by temporal lobe atrophy. Amplifying the evidence from stroke (Buxbaum et al., 2014; Dresang et al., in press; Watson & Buxbaum, 2015), these findings highlight the interplay between semantics and gesture deficits in PPA, and suggest that clinical assessments that include imitation of meaningful hand postures may be useful in distinguishing svPPA from other PPA variants.

## Footnotes

1 Research from other laboratories has adopted different dichotomies. For example, Schmidt et al (2022), Achilles et al. (2019), and Tessari et al. (2021) focused on the distinction between finger postures versus arm/hand gestures.

2 This study was conducted remotely during the COVID-19 pandemic and thus no research images were collected to characterize brain atrophy at the time of behavioral testing.

